# Mapping QTLs for phenotypic and morpho-physiological traits related to grain yield under late sown conditions conditions in wheat (*Triticum aestivum* L.)

**DOI:** 10.1101/2021.06.17.448834

**Authors:** Yaswant Kumar Pankaj, Lalit Pal, Ragupathi Nagarajan, Kulvinder Singh Gill, Vishnu Kumar, Sonali Sangwan, Sourav Panigrahi, Rajeev Kumar

## Abstract

The elevating temperature makes heat stress one of the major issues for wheat production globally. To elucidate genetic basis and map heat tolerance traits, a set of 166 doubled haploid lines (DHLs) derived from the cross between PBW3438/IC252874 was used. The population was evaluated under Normal sown (NS) and late sown (LS) conditions, by exposing to heat stress during *rabi* season. The canopy temperature (CT) showed positive correlations with grain yield, whereas Soil plant analysis development (SPAD) was not significantly correlated and associated with GY in both the normal and late sown conditions. Composite interval mapping (CIM) identified total 12 Quantitative trait loci (QTLs) *viz*., 2 (Normal sown), 10 (late sown) mapped on linkage groups 1A, 1D, 2B, 2D, 3B, 4D, 5B and 6D, during both the crop seasons 2017-18 and 2018-19. Combining the results of these QTLs revealed a major stable QTL for grain yield (GY) on chromosome 3B with 11.84% to 21.24% explaining phenotypic variance under both sowing conditions. QTL for CT and SPAD was detected on chromosome 1A while QTL for GY on chromosome 3B and 5B. The identified QTLs in the genomic regions could be targeted for genetic improvement and marker assisted selection for heat tolerance in wheat. The tools like SPAD and CT could be exploited to screen the large number of breeding lines.

## Introduction

Wheat (*Triticum aestivum* L.) is one of the most widely grown food grain crops in the world. It contributes 20% dietary calories in the human diet (http://faostat.fao.org). Global warming has a significant effect on its yield reduction. The wheat is prone to heat stress which affects its production at a large scale. The elevated temperature has always a major impact on wheat production. The combined effect of heatwaves with drought is detrimental during the anthesis and grain-filling duration, which are the most vulnerable stage affecting the final yield (Ortiz *et al*. 2008). When the temperature exceeds the optimum value, the rate of photosynthesis in wheat decreases (Sage and Kubien, 2007) due to a reduction in the efficacy of photosystem II (Nash *et al*., 1985). According to Hertel *et al*., 2010 the global wheat productivity has lowered down up to 5% as the temperature rises approximately by 0.13C per decade. By 2020, the south asian countries will see rise in maximum and minimum temperature of 1.54c and 1.08c during rabi season. (Bhusal *et al*., 2017). The central and peninsular region of India faces heat stress and north western region affected by terminal heat stress due to late sowing conditions. (Sharma *et al*., 2014). On the other hand, the southern great plains of America reported temperatures of 32-35c during grain filling duration. The high temperature accounts for phenotypic changes as well. For each degree rise in temperature, it is reduced by 6%. According to (Mason *et al*. 2010), if the wheat grain growth period coincides with high temperatures, it can reduce grain yield by up to 28.3%. Heat stress accelerates the wheat growth phase and shortens the grain filling duration leads to lower yield. It creates alteration in traits like morpho-physiological, agronomic, and yield-related characters. The morpho-physiological characters like canopy temperature and chlorophyll are easy to detect with handheld meters and handy for breeders. These traits have a great correlation with yield under high temperatures. Criteria for the selection of heat-tolerant genotypes must be rapid and cheap in the evaluation and should also allow for the screening of a high number of plants in a short period (Mitra 2001; Araus *et al*. 2002). To overcome this abiotic insult, breeding materials should be utilized which helps in understanding the genotype x environment relationship. The breeding population like doubled haploids, which can be utilized to deliver thermo-tolerant lines. It is an important tool for plant genome mapping. Molecular marker is prerequisite to identify quantitative trait loci (QTL) associated with stress tolerance (Campbell *et al*. 2003; Pinto *et al*. 2010). To find the stability of marker-QTL in different environments and genetic background is a tedious job (Yang *et al*. 2007). But, newer and evolving software programmes has lead us to provide more reliable results and helps us in understanding the complexity between genes and the phenotypic traits. Therefore, several efforts have been undertake to modify this crop to tolerate heat. In view of the facts mentioned above, the present study was undertaken to develop marker-QTL linkage involved in heat stress in the DH population. Morpho-physiological characterization of the doubled haploid lines considering heat stress.To discuss the impact of high-temperature stress on wheat and traits associated with heat tolerance which would help formulate management strategies for wheat yield improvement under high-temperature stress and breeding for high-temperature tolerant varieties.

## Materials and methods

### Plant material

A doubled haploid population, derived from a cross between PBW343 (heat susceptible) and IC252874 (heat tolerant) (Fig 1) was sown in four experiments over two field seasons. The population contained 166 individuals. The plants were planted in alpha lattice design with three replications. Proper agronomic practices and irrigation were provided to avoid yield reduction during the crop cycle. The development of the DHs was done in the Washington State University, Pullman, USA, and evaluation was carried out at the research farm of the Rajendra Prasad Central Agricultural University, Pusa, Bihar, India (25° 57′ 08″ N; 85° 40′ 13″ E). An offseason facility at the Research station of Punjab Agricultural University, Keylong, Himachal Pradesh, India, was utilized for seed multiplication. .

**Fig 1:**
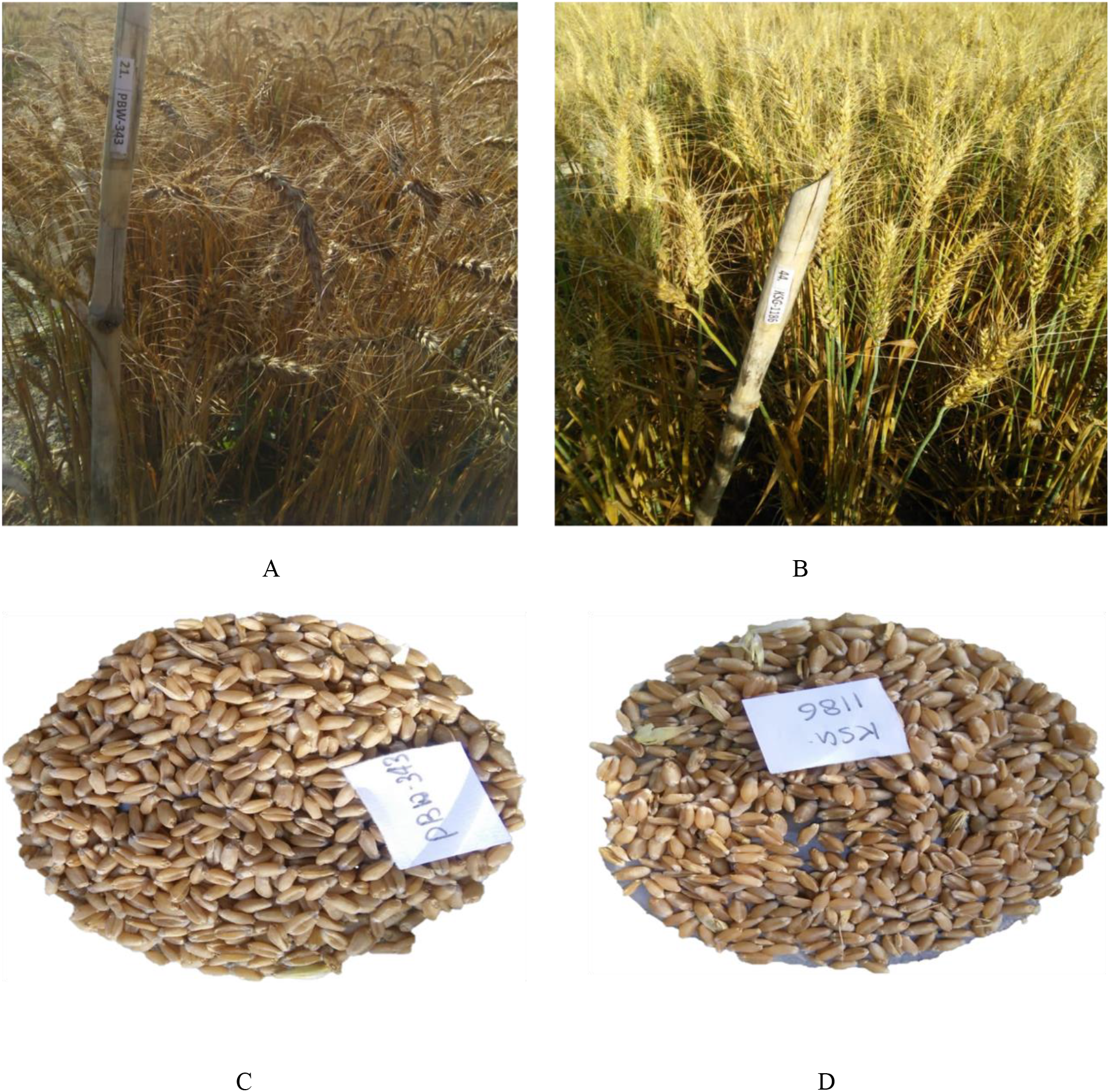
A. PBW343 (Heat susceptible parent) B. IC252874 (Heat Resistant parent) C. PBW343 Seeds D. IC252874.

### Data collection and evaluation for heat stress

To expose the plants to different levels of temperatures at the time of grain filling, the crop was sown during the second week of November and during the first week of January. Then average ambient temperatures during the grain growth phase between anthesis to physiological maturity were 25.5 and 25°C when sown in November and 32 and 32.8.0°C when sown in January in 2017–2018 and 2018–2019, respectively. As differences in days to heading in selected genotypes ranged from 8 to 14 days, previous observations were used for staggered sowing to ensure synchrony in flowering and thus nearly same level and magnitude of exposures to ambient temperatures during grain filling across the genotypes. Irrigation was provided to the crop at five growth stages (at crown root initiation Zadok, GS 21; tiller completion Zadok, GS 29; late jointing Zadok, GS 36; flowering Zadok, GS 61; and milk stage Zadok, GS 75) and six (at crown root initiation Zadok, GS 21; tiller completion Zadok, GS 29; late jointing Zadok, GS 36; flowering Zadok, GS 61; milk Zadok, GS 75; and dough stage Zadok, GS 85) both in timely and late-sown experiments during the crop season in both the years to maintain optimum soil moisture. Weeds were removed manually. To avoid border effects, plants at the centre of each plot were chosen for collecting all the data. The data were recorded for total ten characters viz. heading, was calculated as the length of the period between seedling emergence and the time that 50% of spikes emerged from leaf sheath, maturity was measured with 75% yellowness, plant height and peduncle length was measured on a random sample of five plants in each plot as the distance from the soil surface to the tip of the spike awns excluded at harvest time (cm). Total leaf chlorophyll content (SPAD index) estimated using spad-502 chlorophyll meter (spad-502 plus, Konica Minolta, Kearney, NE, USA), during the flowering stage. Canopy temperatures were measured using a handheld infrared thermometer (KM 843, Comark Ltd., Hertfordshire, UK) with a field view of 100 mm to 1000 mm. Canopy temperatures data were taken from the same side of each plot at 1m distance from the edge and approximately 50 cm above the canopy. CT values were recorded at three growth stages (GS55, GS65, and GS83). Plants at the centre of each plot were selected to count the number of tillers at the growth stage, GS85; (Zadoks *et al*. 1974). Both shoots and roots were collected for biomass determination. Readings were made between 1300 and 1500 h on sunny days. Thousand-grain weight (TGW in g) and grain yield (GY in g) was calculated by weighing grains of harvested plants from an area of 2 m^2^ in each plot excluding border effect (Sayre *et al*. 1997).

### Statistical analysis

Analysis of variance and least square means of all traits were estimated using the statistical procedure Proc. Mixed of SAS version 8.2 (SAS Inst. Inc. 1999).Values of means, medians, standard deviation, Variance, Coefficient of variation, minimum, maximum, and range showing the distribution of phenotypic data for different traits were determined using R software. Correlation coefficients between the traits for the trials were performed using R software. A stepwise regression analysis-based clustering was performed. The differences in trait response between the normal and stress conditions for phenotypic factors corresponding under stress were considered to perform cluster analysis using pair group distance with Euclidean distance measures. The dendrogram was constructed using the R software (version 3.5.1).

### Genotyping, construction of linkage map, and QTL mapping

Few seeds of parents and DHs were germinated on filter paper in dark and coleoptiles were used to extract DNA following the recommended DNA extraction method (http://www.triticarte.com.au/content/DNA-preparation.html. The thermocycling programme consisted of an initial denaturation at 94°C for 3 min, followed by 30 cycles of 30 s at 94°C, 30 s at 50/65°C, 30 s at 72°C and a final cycle of 2 min at 72°C in Thermal Cycler (Sharma *et al*. 2016). A total of 200 SSR markers were screened for polymorphism between the parents and the resulting polymorphic markers were used to screen the DHs. Profiles of polymorphic SSR markers were scored visually by coding PBW343 alleles as “A” whereas IC252874 alleles were scored as “B”. While heterozygote individuals are scored as “H”. Missing bands were scored as ‘NA’.A consensus map (Somers *et al*. 2004) was used to select microsatellite/simple sequence repeat. Allele bands were visualized on 2% agarose gels. (Fig2) For the construction of linkage maps, a set of 61 SSRs (gwm, wmc, swm, barc, and cfd) spanning on eight wheat chromosomes (1A, 1D, 2B, 2D, 3B, 4D, 5B, and 6D) were deployed during the present investigation. Out of 200 SSR markers, 12 markers were not amplified at all and 61 SSR markers that were polymorphic between the parental genotypes of the DH mapping population were used for the preparation of the linkage map. The genotyped data of the DH population was used to generate a linkage map using software MapDisto 2.1.7.1.

**Fig 2:**
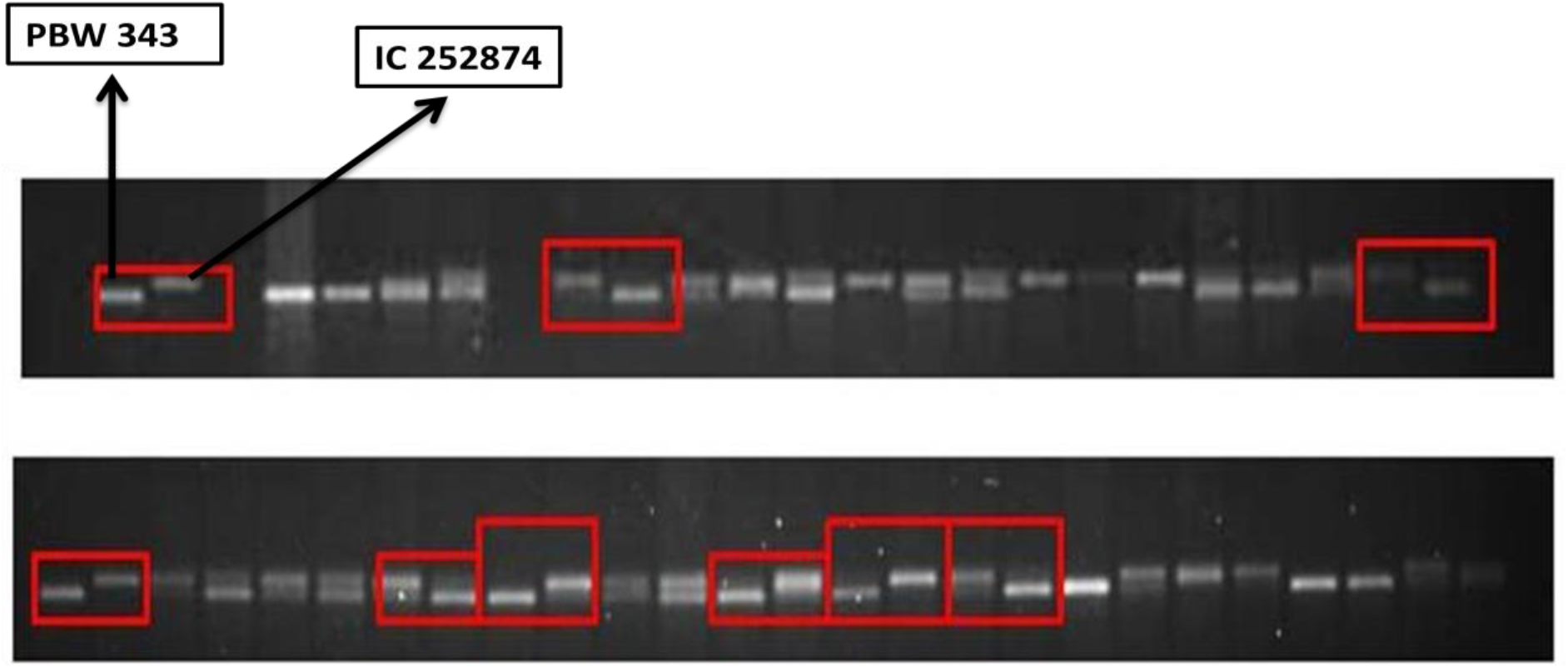
Parental polymorphism between PBW343 and IC252874.

The Kosambi mapping function and interval position type was used for the conversion of recombination frequency into the genetic distance. QTL analysis was performed using QTL Cartographer v2.5 (Wang *et al*., 2010). In the CIM method, forward regression with five background markers, a window size of 10.0 cM, and a walking speed of 2 cM were used in the software Windows QTL Cartographer 2.5 (Wang *et al.* 2010). The trait setting for Composite interval mapping (CIM) was done using model 6 and used to determine likely QTL positions and a threshold of 1,000 permutation test at P = 0.05. The putative QTLs were defined as two or more linked markers associated with a trait at LOD > 3.0. The suggestive QTLs were defined as QTLs where two or more linked markers were detected at 2.0 < LOD < 3.0 (McIntyre *et al*. 2010). Here, The LOD value was set as minimum LOD value 2.0, to keep consider suggestive and minor QTL as well.

## Result

### Phenotypic assessment for heat tolerance

Very contrasting differences were found in parental lines for plant height and grain yield under stress condition. The heat-tolerant parent, IC252874, showed non-significant reduction for all yield components, whereas plant height was found highly reduced (20% and 19.2%) under late-sown condition in both the years. On the contrary, the heat-sensitive parent, PBW 343, showed significant reduction in plant height (24% and 21%), CT (27% and 29%) and GY (40% and 33%) with little reduction in SPAD under late-sown condition in both the years. The heat-tolerant genotype IC252874 showed an increase in the DM with a shorter growth period, whereas the heat-sensitive genotype PBW 343 showed medium days to maturity. It was interesting that there was no significant reduction in TGW in both parents under terminal heat stress in late-sown condition as compared to timely sown. A significant reduction in DM (18% and 19.2%), CT (19% and 21%), PH (16% and 14.3%) and GY (33% and 30.1%) was observed in DH population under heat stress across the years. Mean values were normally distributed within the DH population due to the presence of a large number of transgressive segregants (Table 1). The doubled haploid lines along with the parents were subjected to genetic divergence analysis differed significantly with regard to the above studied characters and displayed marked divergence and grouped into eight clusters following Euclidean method (Table). Cluster I is the largest cluster with sixty one DH lines and a parent, PBW 343 viz. DH 61, DH 60, DH 45, DH 48, DH 50, DH 53, DH 54, DH 57, DH 63, DH 64, DH 67, DH 73, DH 74, DH 78, DH 80, DH 89, DH 91, DH 112, DH 120, DH 123, DH 127, DH 128, DH 133, DH 134, DH 135, DH 138, DH 142, DH 145, DH 146, DH 152, DH 159, DH 160, DH 161, DH 163, DH 164, DH 166, DH 24, DH 25, DH 26, DH 27, DH 28, DH 29, DH 31, DH 32, DH 34, DH 35, DH 36, DH 38, DH 39, DH 43, DH 1, DH 6,PBW, DH 8, DH 9, DH 10, DH 12, DH 13, DH 15, DH 17, DH 59, DH 72 followed by Cluster VI having thirty four DH lines namely DH 95, DH 121, DH 124, DH 100, DH 49, DH 56, DH 141, DH 58, DH 139, DH 33, DH 65, DH 69, DH 157, DH 52,DH 155, DH 85, DH 23, DH 46, DH 68, DH 66, DH 86, DH 84, DH 158, DH 14, DH 20, DH 21, DH 30, DH 83, DH 147,DH 153, DH 154, DH 22, DH 55, DH 75. Cluster V, and VIII has twenty five, seventeen and twenty one DH lines while Cluster II, III and IV accounted for five, eight and one DH lines. (Fig 3)

**Table 1:** Mean, maximum and minimum of clusters using phenotypic data.

| Cluster | Parameters | DHE | DM | PH | TGW | GY | CT | SPAD | BMAS |
| --- | --- | --- | --- | --- | --- | --- | --- | --- | --- |
| I | Mean | 71.322 | 109.47 | 98.46 | 36.45 | 145.61 | 35.09 | 42.11 | 466.2 |
|  | Max | 61 | 100 | 54.6 | 19.06 | 95 | 15.98 | 19.18 | 98 |
|  | Min | 79 | 116 | 137 | 81.2 | 223 | 61.99 | 74.39 | 674.5 |
| II | Mean | 68.6 | 111.2 | 100.94 | 37.8 | 141.2 | 33.49 | 40.19 | 295 |
|  | Max | 65 | 108 | 81.6 | 19.1 | 95 | 17 | 20.4 | 220 |
|  | Min | 70 | 113 | 116 | 80.1 | 221 | 59.99 | 71.99 | 500 |
| III | Mean | 71.25 | 109 | 103.55 | 31.69 | 136.41 | 32.2 | 38.64 | 221.5 |
|  | Max | 66 | 101 | 57.8 | 19.98 | 97.3 | 15.98 | 19.18 | 101 |
|  | Min | 76 | 113 | 131 | 41.02 | 169 | 45.01 | 54.01 | 525 |
| IV | Mean | 73 | 111 | 94 | 77.02 | 205 | 57.1 | 68.52 | 441.1 |
|  | Max | 69 | 120 | 90.01 | 82.03 | 201 | 63.2 | 75.3 | 340 |
|  | Min | 78 | 128 | 100.2 | 100.23 | 194.5 | 56 | 62 | 234 |
| V | Mean | 72.375 | 109.46 | 104 | 31.26 | 134.67 | 32.39 | 38.87 | 505 |
|  | Max | 63 | 104 | 65 | 19.23 | 94.8 | 15.7 | 18.84 | 101 |
|  | Min | 81 | 116 | 142 | 45.09 | 183.3 | 55.18 | 66.22 | 735 |
| VI | Mean | 71.059 | 109.59 | 95.74 | 31.98 | 137.35 | 32.3 | 38.76 | 403.2 |
|  | Max | 65 | 102 | 58 | 20.02 | 95 | 17.2 | 20.64 | 250 |
|  | Min | 77 | 115 | 131 | 42.09 | 177 | 50.09 | 60.11 | 685 |
| VII | Mean | 60 | 100.01 | 80.2 | 28.65 | 89.05 | 23.45 | 30 | 421.95 |
|  | Max | 71.29 | 108.53 | 92.79 | 35.88 | 145.54 | 36.05 | 43.26 | 645.2 |
|  | Min | 64 | 101 | 63.5 | 20.12 | 97 | 16.34 | 19.61 | 500 |
| VIII | Mean | 60.54 | 100 | 95 | 30 | 85 | 23.64 | 32.21 | 300.53 |
|  | Max | 70.857 | 108.38 | 102.41 | 38.51 | 153.82 | 37.5 | 45 | 531.3 |
|  | Min | 66 | 104 | 75 | 20.98 | 98 | 18.22 | 21.86 | 280 |

**Fig 3:**
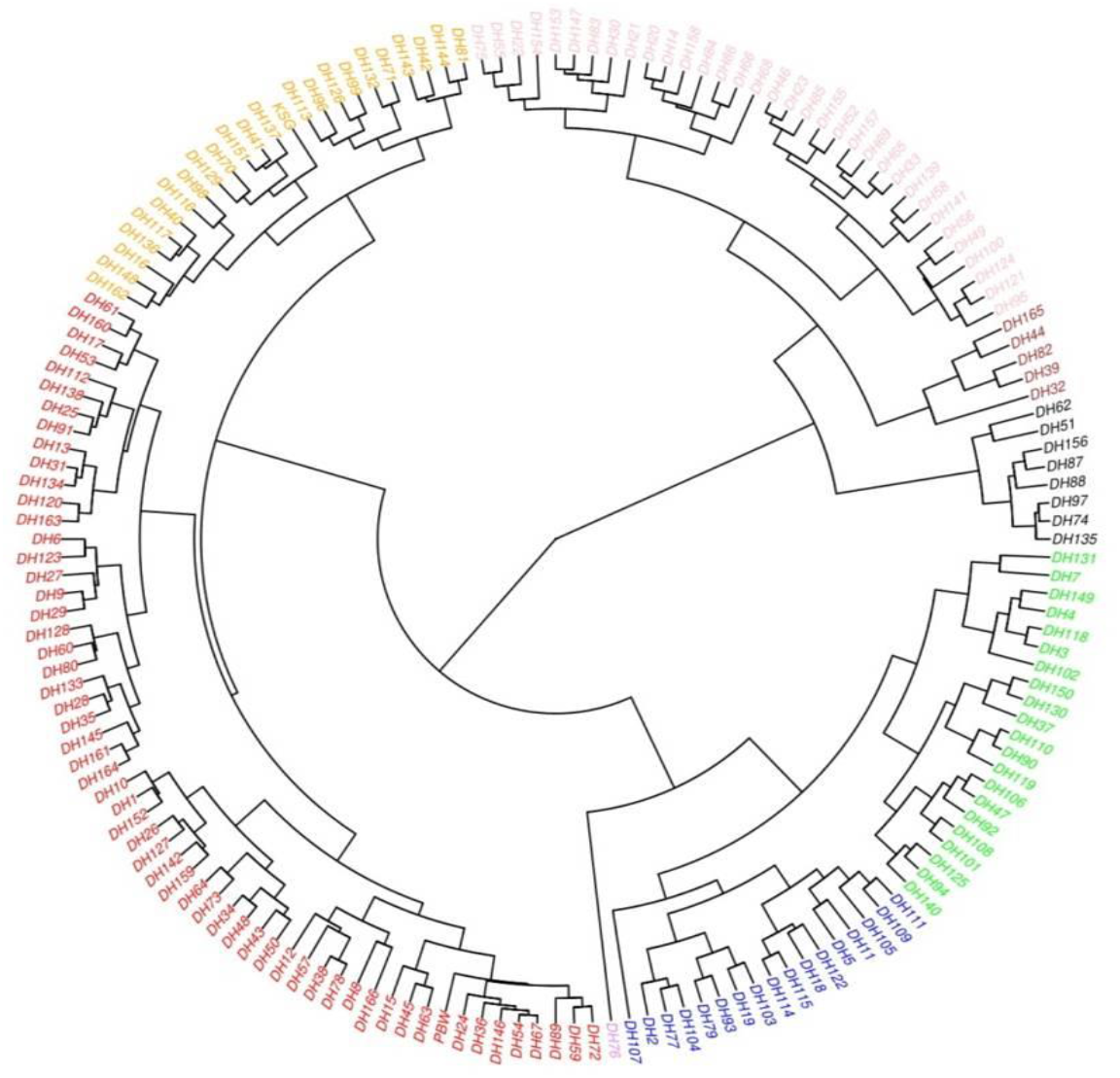
Dendrogram on the basis of phenotypic diversity present in doubled haploid lines.

### Weather condition and phenotypic summary

The late-sown heat experiments were the hottest in terms of average minimum and maximum temperatures during vegetative growth and grain fill, and had the highest number of days with temperature (28 and 34C). The meteorological data for both the cropping season 2017-2018 and, 2018-2019 is given in (Table 2). Analysis of variance showed significant genotypic variance present in the DHs for heat tolerance (Table 3). For the eight studied traits, values of correlation coefficients based on data pooled over Normal sowing and Late sowing conditions are presented in (Figure 4). Each of the traits showed a positive and significant correlation across year, confirming the stability of these measurements of high temperature tolerance. Both in the NS and LS conditions, CT was non-significantly correlated to DH, The magnitudes of correlation of CT with other traits were higher under LS condition relative to those in the NS condition. The correlation of GY with TGW was positive and significant, while with CT, it was negative. The correlation of SPAD with GY and TGW was low to moderate.

**Table 2:** Peak temperature, average maximum and average minimum temperature, rain, relative humidity and pan evaporation during the cropping seasons (2017–18 and 2018–19)

| Year | Trial | Peak Temp (month) (oC) | Sowing date | Harvesting date | AvgMax/AvgMin (oC) | Rain (mm) | RH (%) |
| --- | --- | --- | --- | --- | --- | --- | --- |
| 2017-18 | Control | 31 (Apr) | 22 Nov | 11 Apr | 25.2/11.2 | 18 | 46.5 |
|  | Late sowing | 39 (May) | 21 Jan | 19 May | 33.3/15.3 | 16 | 40.3 |
| 2018-19 | Control | 30 (Apr) | 22Nov | 11Apr | 23.2/12.1 | 19 | 43.8 |
|  | Late sowing | 38 (May) | 21 Jan | 29May | 31.1/12.3 | 20 | 37.9 |

**Table 3:** Pooled Analysis of variance for different morpho-physiological and yield traits.

| Source of Variation | df | DHE | DM | PHE | TGW | GY | CT | SPAD | BMAS |
| --- | --- | --- | --- | --- | --- | --- | --- | --- | --- |
| Replications | 3 | 36.04** | 40.5** | 0.0087** | 10.60 | 12483.77 | 2.64* | 9.7** | 4120** |
| Year | 2 | 8.6* | 854.1** | 639.9** | 3553.56** | 1476872.00** | 3770.45** | 7047.3** | 428 |
| Line | 166 | 72.6** | 40.5** | 0.0143* | 33.59** | 293175.25** | 1.75** | 2.8** | 350 |
| Year X Line | 322 | 24.1** | 19.0** | 76.8** | 23.10** | 278252.75** | 1.55** | 1.97 | 6513** |
| Error | 762 | 14.27 | 33.09* | 4.179 | 11.10 | 157771.44** | 0.73 | 0.84 | 110.98* |
| Heritability |  | 0.67 | 0.53 | 0.50 | 0.67 | 0.72 | 0.70 | 0.68 | 0.50 |

**Fig 4:**
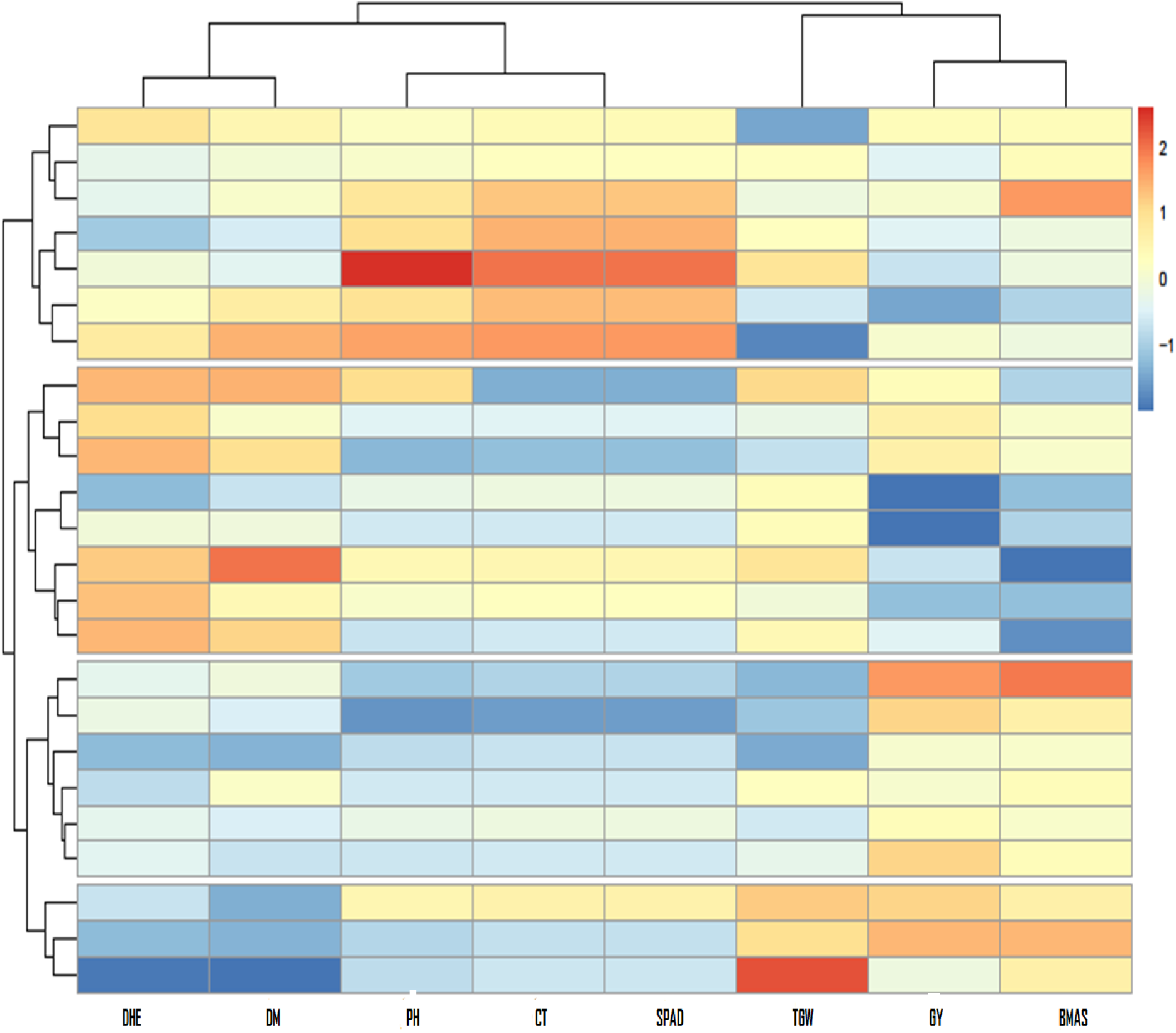
Heat maps illustrating the phenotypic correlation between all traits measured in the doubled haploid population PBW343/IC252874.

### Molecular characterization of DH population and genetic linkage map construction

A set of 200 SSR primers were used to detect polymorphism between parental genotypes, out of Which, 61 SSRs were polymorphic. The sixty-one SSRs showed a good fit to the 1:1 segregation ratio in the DH mapping population. The linkage maps spanned 3,314.64 cM coverage. The maximum number of polymorphic markers were found on 2B with fourteen markers while lowest being two on 6D chromosome. The linkage map developed using 61 polymorphic markers and 166 DHs population was further utilized to analyze QTLs. The genome of DHs is composed of 61per cent of PBW 343 whereas 23.4 per cent of IC and remaining is 9 percent, constitute the heterozygous, distorted and missing values. The linkage maps with identified QTLs in PBW343/IC252874 DHs were presented in (Fig. 5)

**Fig 5:**
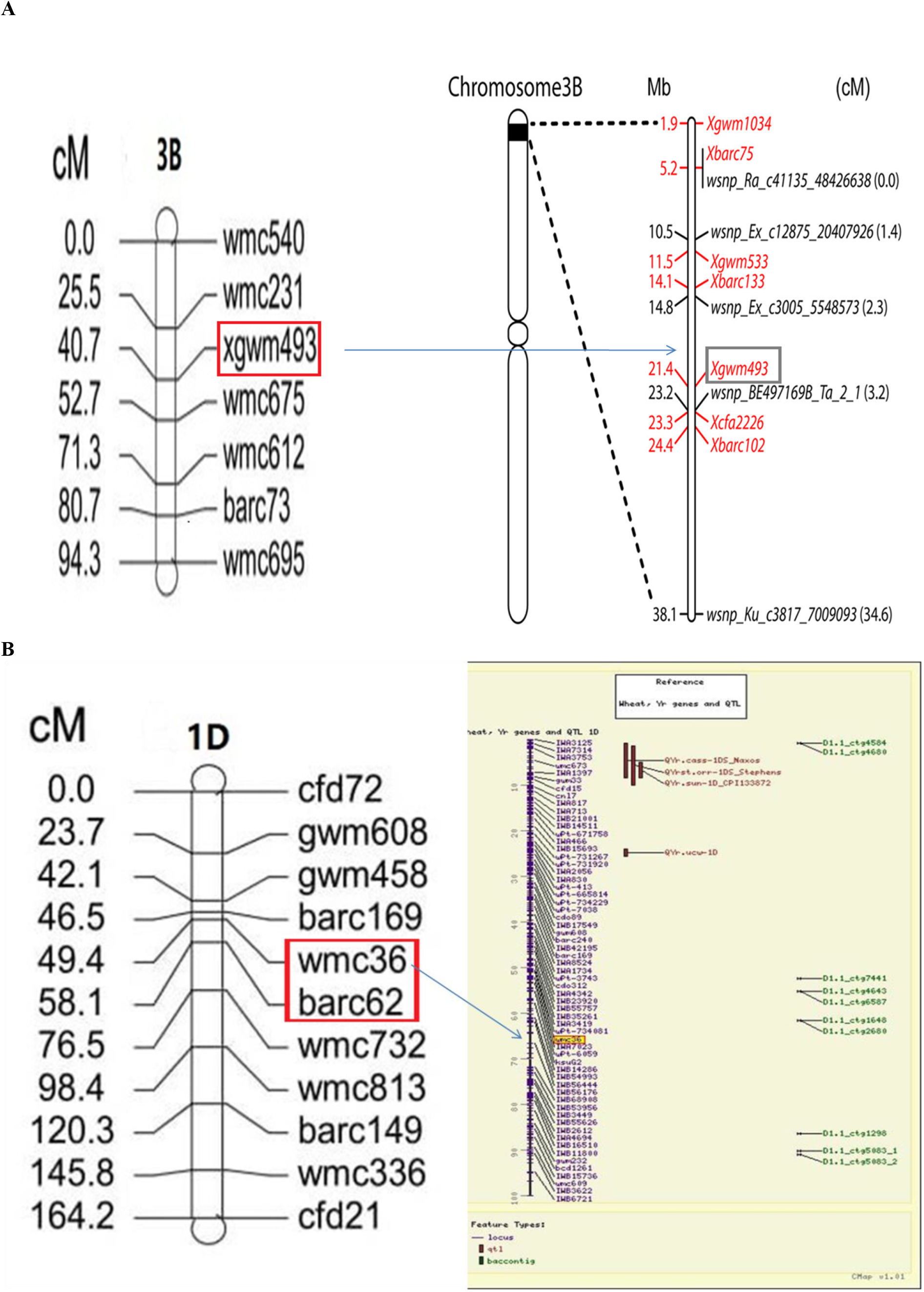
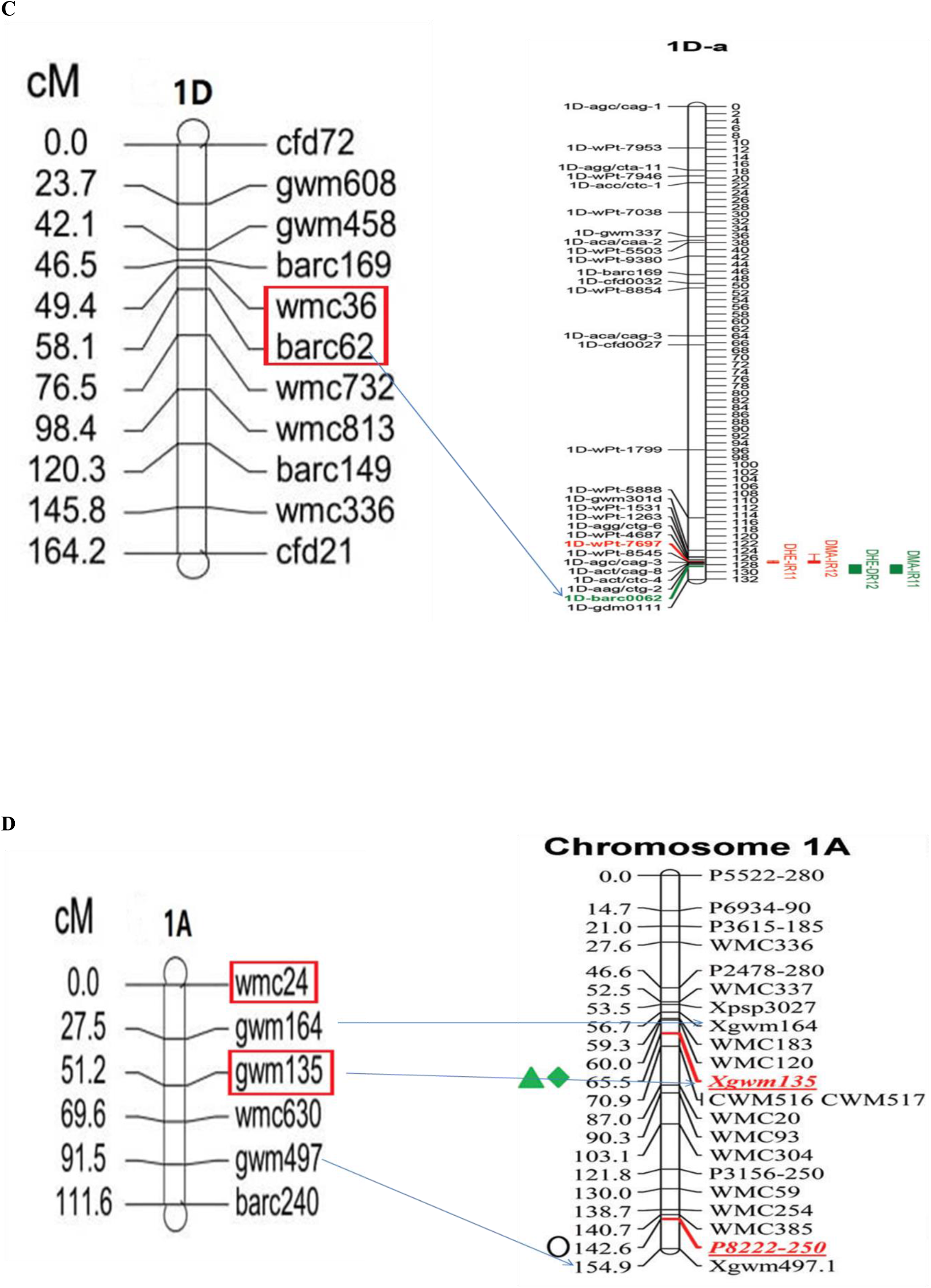
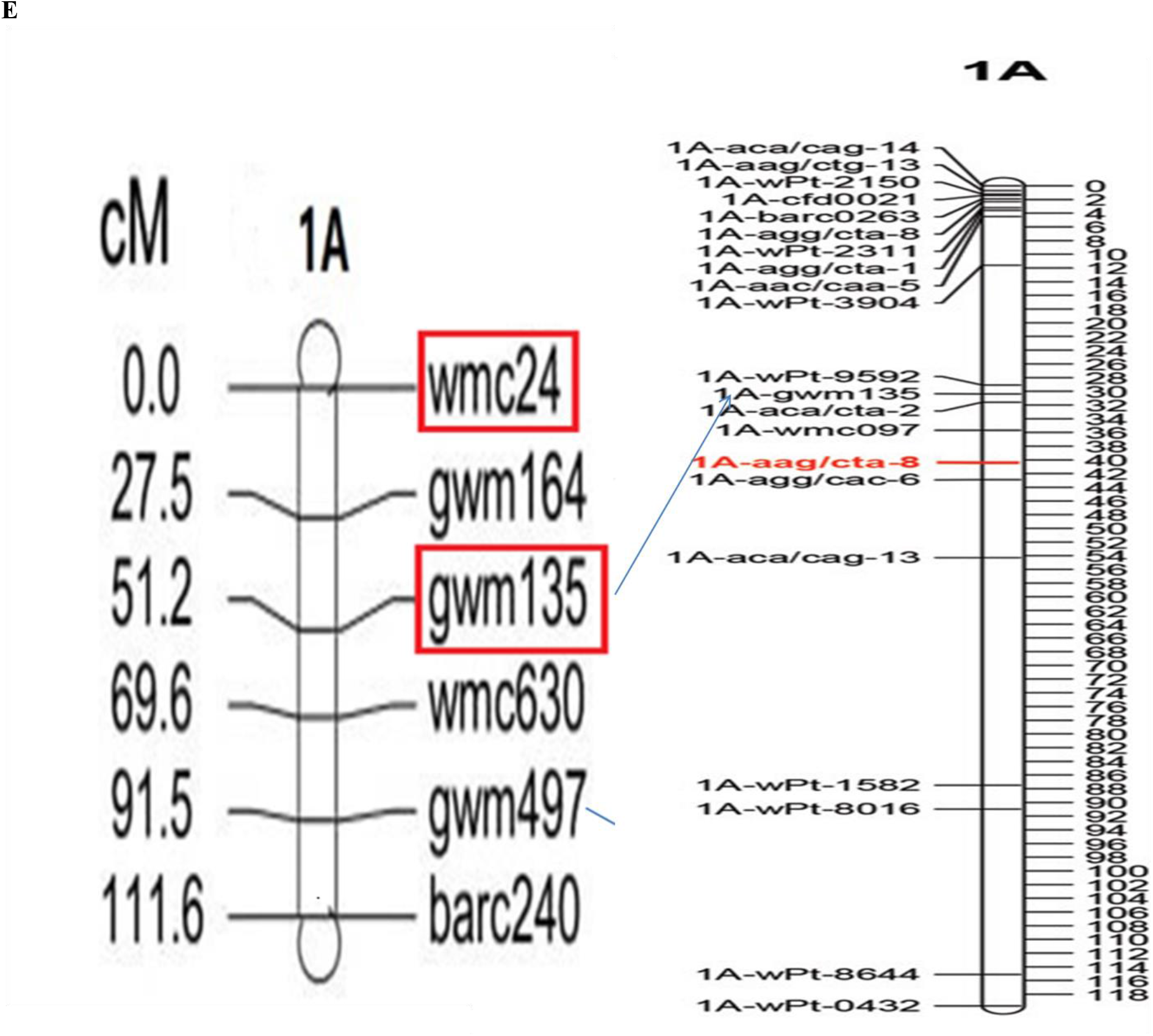
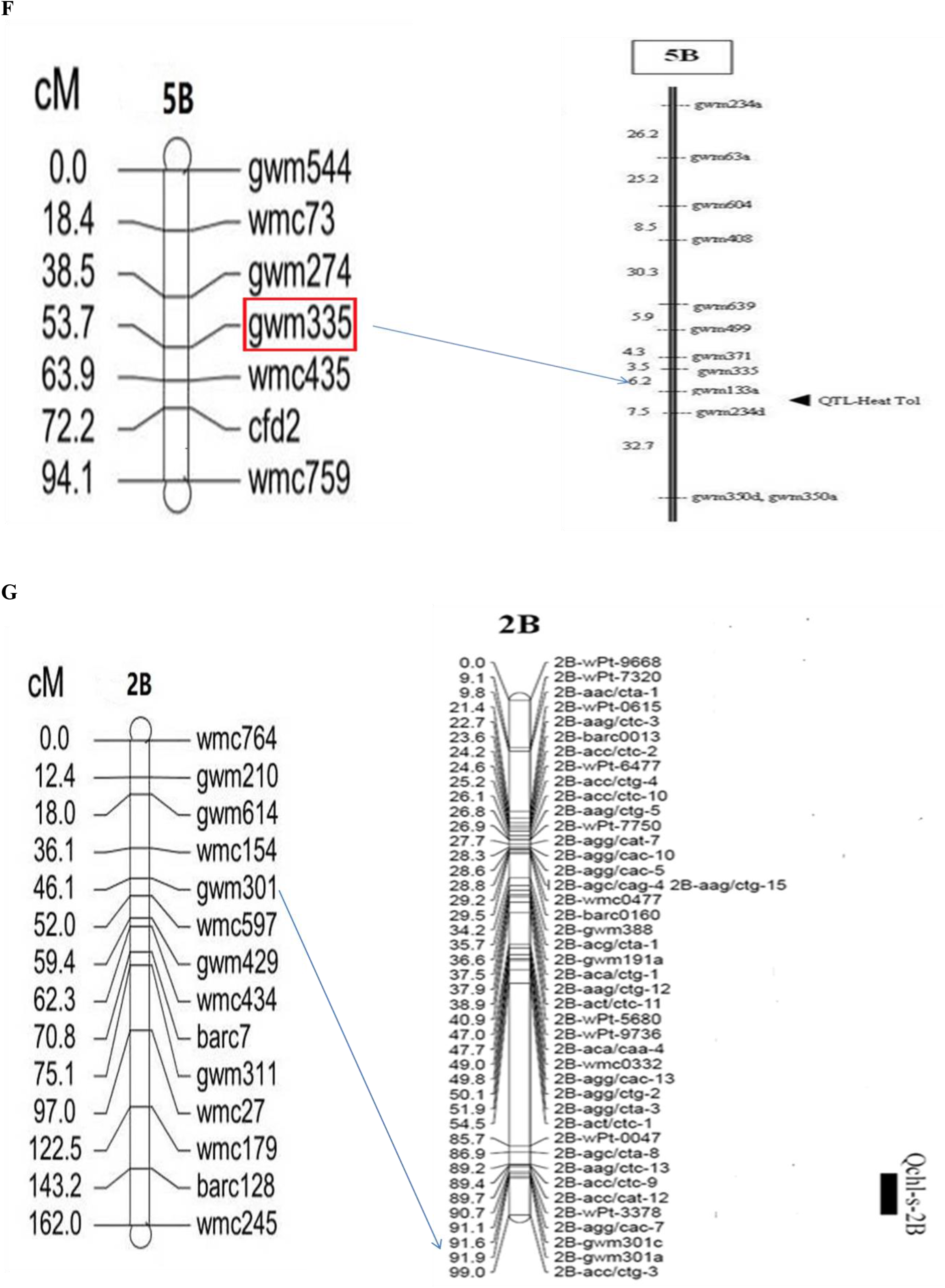
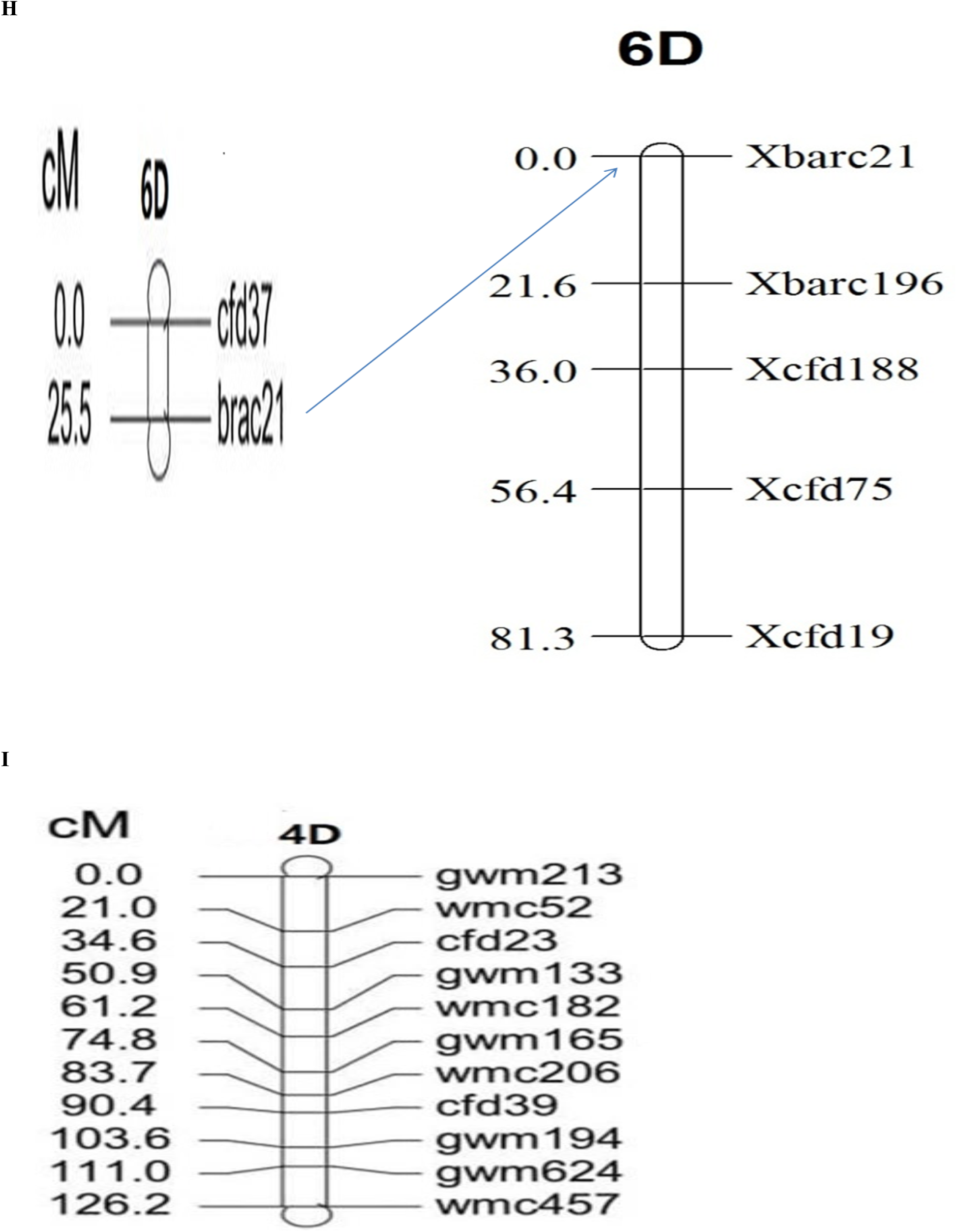
Genetic linkage map (A, B, C, D, E, F, G, H, I) of PBW343/IC252874 DH population with identified QTLs.

### QTL mapping for the heat stress tolerance

A total of 12 QTL were detected for eight different traits using Composite Interval Mapping (CIM) under Normal sown and late sown condition during both the crop seasons 2017-2018 and 2018-19 (Table 4). These were located on 8 different chromosomes, 1A, 1D, 2B, 2D, 3B, 4D, 5B, and 6D. The LOD scores of identified QTLs varied from 3.02 (DM-4D) to 21.35 (CT-1A) during 2017-18 and 2018-19 explaining 6.72% and 25.61% phenotypic variance, respectively (Fig 6). A minimum of 1 QTL were available each for, DHE, TGW and CT, 3 QTLs were available for DM and SPAD on an average. A maximum of 4 QTLs were detected for GY under Normal and late sown conditions, respectively. QTLs were identified both in normal sown (2 QTLs) and late sown conditions (10 QTLs). In this study, QTLs for biomass and plant height were not found either in normal or late-sown conditions across the year. The absence of QTLs for phenological traits indicated that there were no confounding effects between environment and phasic development.

**Table 4:**
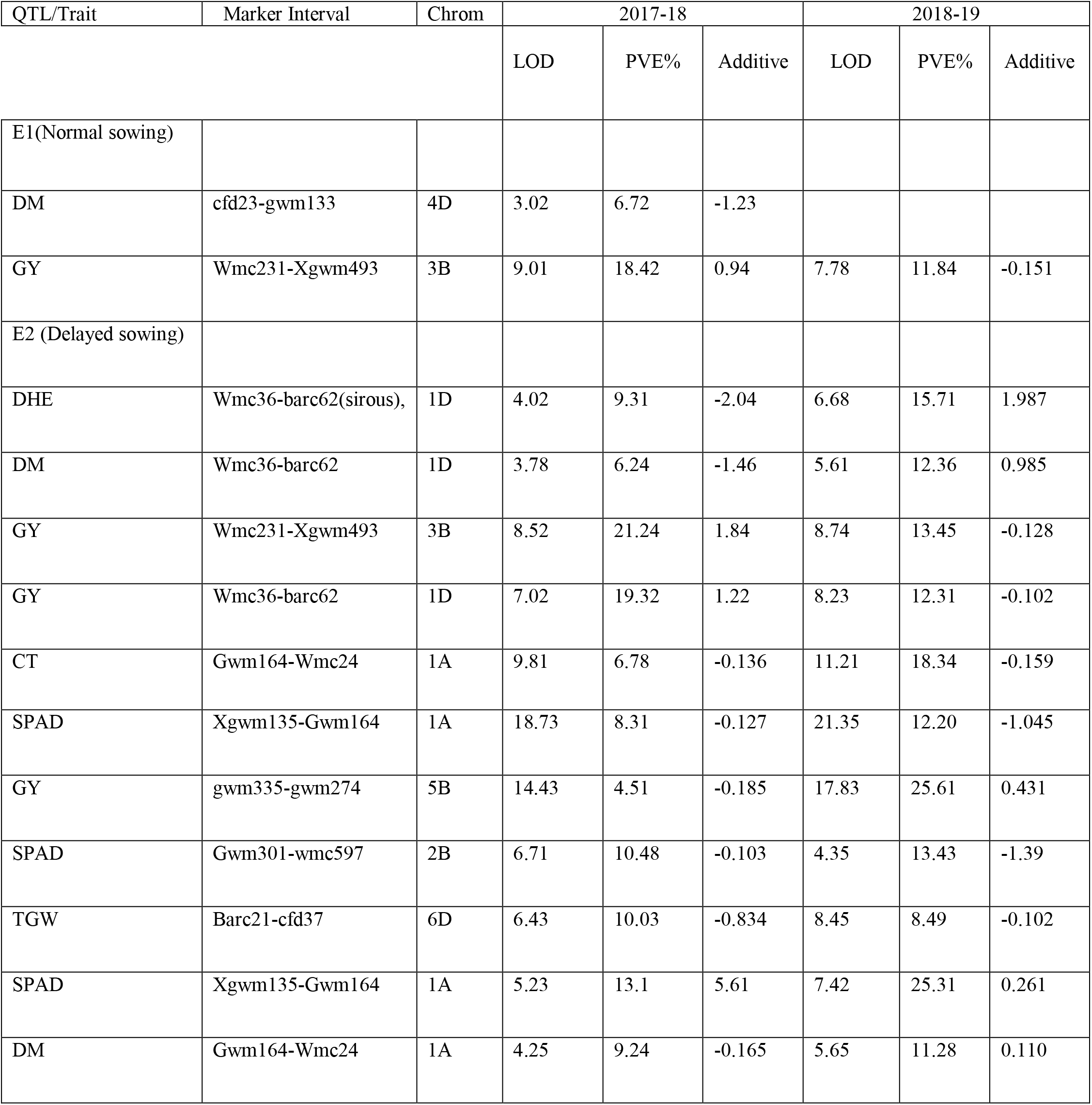
NS (Normal sown) and LS (late sown), Position LOD, A LOD threshold of 2.5 was used for declaration of QTL, PVE (%), Phenotypic variance explained by QTL, Additive effect, Positive “additive effect “indicates an increasing effect from PBW343; negative “additive effect” indicates an increasing effect from IC252874.

**Fig 6:**
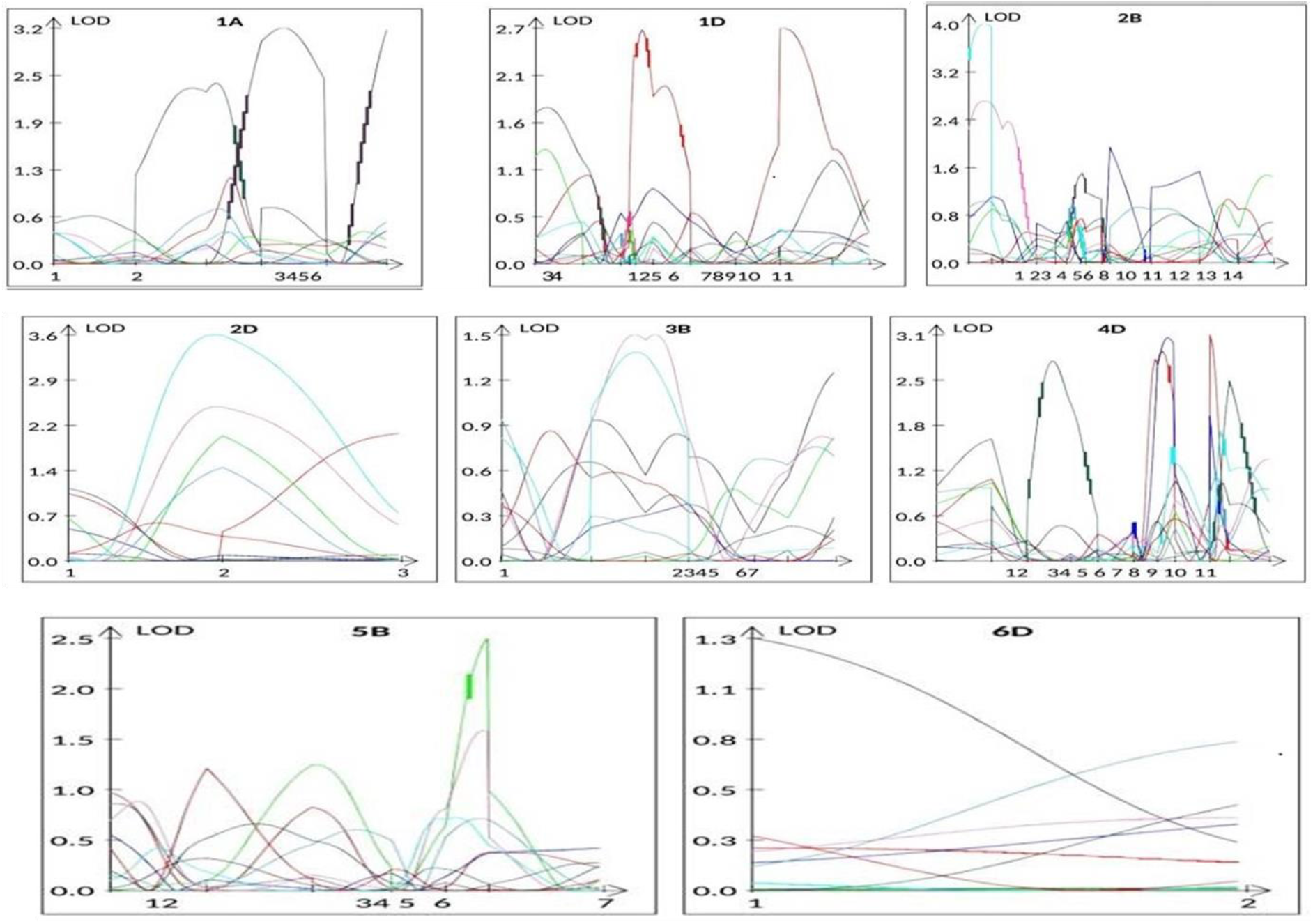
Representative QTL hits detected for various physiological and yield component traits in the PBW343/IC252874 DH population.

## Discussion

To combat the climatic insult in agriculture, molecular resources are used to develop stress resilient crops. QTL mapping is a best known approach to unravel the molecular information linked with the plants phenotype under various environmental (stress) conditions. Several QTL mapping studies in wheat under abiotic stress conditions involved mapping populations such as RILs (Pinto *et al*. 2010; Mason *et al.* 2011; Paliwal *et al*. 2012), doubled haploids (Tiwari *et al*. 2013) and segregating F2 and F3 populations (Gupta *et al*. 2015). In present investigation, a doubled haploid population is used to undertake this study. The ability to manage different sowing conditions created two distinct environmental treatments for the PBW343/IC252874 doubled haploid population. The phenotypic response to normal and late sown conditions within the population has been unraveled by QTL mapping. The late sown condition was the lowest yielding on an average for the population and it resulted into shortest ear emergence time, caused due to higher temperature and longer day length. Due to the longer growing season under late sown condition, it was warmer. During flowering time, it was more prone to abiotic stress such as high temperature (Dolferus *et al*. 2011; Gibson and Paulsen 1999). There was a significant variation present in the parents of the DH population for the studied traits suggesting genetic diversity among both the parents. Analysis of variance showed significant (p < 0.01) main effects due to genotypes for all studied traits present in the DHs for heat tolerance. The frequency distribution of DHs exceeded beyond those of the parents reveals the presence of transgressive segregation. As observed by Yang *et al*. 2002, this suggested that the parents contributed different genes for heat tolerance and that traits were not simply inherited. Rieseberg *et al*. 1999 suggested complementary gene action as a primary source of transgression and Bhusal *et al*., 2017 observed the RILs population also showed transgressive segregation; exceeding both the parents for traits indicated genes with positive and negative effects were dispersed between parents.

According to the above weather forecast, the late sown condition showed terminal heat stress towards the crop over both the year. The mean minimum and maximum temperatures, under late sown condition were higher than the normal sown condition during 2017-2018 and 2018-2019 crop seasons. However, heat stress during 2018-2019 crop seasons was comparatively more than 2017-2018.

Phenotypic correlation between the traits explains that DHE was negatively correlated with GY under late sown condition.The correlations obtained in present study are in agreement with the previous reports under heat stress. Days to maturity was significantly correlated with, TGW, GY under control and late-sown conditions. Phenotypic correlations of TGW with YLD were significant and positive (0.15–0.89) (Tahmasebi *et al*. 2016). Both DHE and DMA had negative and significant genotypic correlations with GY. Negative phenotypic correlations were found between DHE and TGW. PHE showed positive genetic correlations with DHE, DMA, but its correlations with CT, was negative. Themorpho-physiological characters such as canopy temperature (Bahar *et al*., 2008), and cholorophyll content (SPAD) (Yıldırım et al., 2011) provides genetic gain to wheat. To characterize the mapping populations and genotypes under heat stress condition, CT has been widely used (Tiwari et al. 2012; Reynolds et al. 1997). In this study, the tolerant parent displayed higher value for CT in late sown conditions which indicated a better cooling capacity during grain filling under higher temperature. Similarly, reported by Reynolds *et al*. 1994 and Ayeneh *et al*. 2002. The degree of cooling reflects the rate of evapotranspiration on the surface of the plant canopy (Ayeneh *et al*. 2002) and gains in yield owing to the positive effect of reduced canopy temperature (Reynolds *et al*. 2007). Ayeneh *et al*. 2002 reported that canopy temperature can be used as a tool in the selection of wheat targeted for tolerance to heat stress. We have found positive correlations of canopy temperature with grain yield. According to Reynolds *et al*., 1994, Fischer *et al*., 1998, Phenotypic correlations of CT with grain yield were occasionally positive. However, Saint Pierre *et al*. 2010 reported significant negative phenotypic correlation (r = −0.34 to −0.75, P\0.001) between canopy temperature and grain yield under drought conditions in wheat. On the other hand, SPAD index was not significantly correlated and associated with GY in both the normal and late sown conditions. Similarly, Tahmasebi *et al*. 2016 reported that there was no consistent association between SPAD and GY under different environments. On the other hand, Pinto *et al*. 2010 observed, tolerance to heat in field crops, including bread wheat, is associated with a variety of physiological, biochemical, and morphological traits. In this study, three hundred and two hundred SSR primer pairs representing eight chromosomes of wheat were used to detect polymorphism between the parental genotypes, IC252874 (heat tolerant) and PBW343 (heat susceptible). Out of the 200 SSR primer pairs, polymorphism between the parents was detected by 61 (17.3 %) SSRs. Similarly, Pandey *et al*. 2013 screened the parent, Raj 4014, and WH 730 with three hundred SSR markers. Out of those SSR markers tested, 15% were found polymorphic and Bhusal *et al*., 2017 used three hundred and eighty SSR primer, screened with parents HD2808 and HUW510 to detect polymorphism. Of these, 14.2% of markers revealed parental polymorphism. In this present study, QTL analysis allowed mapping of as many as 12 QTLs (NS and LS conditions) for morpho-physiological and yield traits suggesting that the genetic control of these traits in wheat is complex. This was also supported by the continuous distribution in the component traits of heat tolerance. The findings of earlier studies for identification of QTL also suggested complex genetic control of yield and related traits (McIntyre et al. 2010; Pinto et al. 2010; Bennett et al. 2012a; Bennett et al. 2012b; Kadam et al. 2012; Lopes et al. 2012). Use of morpho-physiological traits and tools like CT and SPAD in combination with QTL mapping provides basic understanding and can serve as useful criterion in selection of heat tolerant genotypes (Harikrishna *et al*., 2020).

Composite interval mapping revealed genomic regions on chromosomes 3B harbored a major QTL for GY, will be vital for genetic improvement perspective of wheat for heat stress using marker assisted selection from the yield point of view. This QTL was detected both under NS and LS conditions, suggesting this QTL is more stable and can be used for selection of yielding genotypes for testing under both conditions. This QTL explained 11.84% to 21.24% of phenotypic variance under NS and LS conditions. However, Other published QTL effects were, Grain yield and plant height in stressed and other environments in a durum wheat Kofa × Svevo RIL population, was also associated with Xgwm493 (Maccaferri *et al*., 2008). The chromosome no. 5B also carried a QTL for GY and the marker associated was gwm335. Mohammadi *et al*., 2008 reported a marker gwm133 directly linked to heat tolerance whereas gwm335 was only 6.2cM away from it. So, gwm335 could be useful in carrying heat tolerance gene as well. QTLs for days to heading, days to maturity and grain yield under late sown condition were also detected on chromosome 1D. The LOD score of these QTLs was above 3.0 while, the phenotypic variations of these QTLs were varied from 6.24 to 19.32%. Similarly, Tahmasebi *et al*. 2016 found that 1D-barc0062 was linked to QTL for DHE and DMA. The M-QTL located on chromosome 1D explained 23.7% of DHE variation, respectively. Zikhali *et al*. 2014 recently validated a major Eps QTL in the barc62 marker of 1D using wheat near isogenic lines (NILs). In our study, the QTL identified at the same marker possibly is related to the Eps genes. One QTL was found to be localized on 2B chromosome for SPAD, accounted for a phenotypic variance of 10.48-13.43%. Similarly, Hassan *et al*., 2018 reported two QTLs associated with GY and SPAD on chromosome 2B were also co-located. These common genomic regions for different traits can be explained by the linkage between two or more genes or the pleiotropic effect of one gene. A QTL for TGW was found on 6D chromosome under late sown condition and showed a phenotypic variance of 6.43%-8.45% Similarly, Bhusal *et al*., 2017 also reported a QTL for TGW, associated with barc21, which explained a phenotypic variance of 13.24% but under non-stressed condition. It was interesting to note that the QTLs detected under heat-stressed conditions for SPAD on chromosome 1A also co-localized with the QTL for CT under late sown conditions. Co localization of QTLs is common in wheat and is the hot spot for networking of complex traits such as yield. The significance of this has been reported for heat stress-related traits in wheat (Mason *et al*. 2010, Pinto *et al*. 2010, Paliwal *et al*. 2012). The clusters of QTLs in a few genomic regions increase the validity and utility value of the detected QTLs. In the present study, common genomic regions found for CT and SPAD variations indicated that a group of linked and co-located QTLs on chromosome 1A and affected phonological and yield-related traits. This stability in QTL across the year could be attributed to high stability for these traits and precision in phenotyping. The authors realize that it is important to observe the performance of this population with more multi-location trial and subject it to the SNP platform for the sake of accuracy and precision.

## Abbreviation

SPAD: Soil plant analysis development
CT: canopy temperature
QTL: Quantitative trait loci
NS: Normal sown
LS: late sown
DH: Doubled haploid
GY: Grain yield
DHE: Days to heading
DMA: Days to maturity
PH: Plant height
BMAS: Biomass
TGW: Thousand grain weight
CIM: Composite interval mapping

## Declaration

NA

## Acknowledgement/Funding

The work was supported by United States Agency for International Development-Biotechnology Industry Research Assistance Council (Reference number: BIRAC/TG/USAID/08/2014) and Acknowledgement to the Department of Agricultural Biotechnology and Molecular Biology, Rajendra Prasad Central Agricultural University, Pusa, Bihar

